## Supplementary material for "Fish-microbe systems in the hostile but highly biodiverse Amazonian blackwaters": Suppl. Mat.

† in memorian

#### Ecology of the sampled Amazonian fish species

The four wild Amazonian host species collected for this study were chosen due to (1) their large-scale distribution in the Amazonian basin, (2) their relatively high abundance at every sampling site, making it possible to collect 20 specimens per species per site, and (3) their tolerance to the contrasting environmental conditions (three Amazonian water types) (Van der Sleen and Albert, 2018). The species selected included two Cichliformes (*M. festivus* and *Cichla spp*.) and two Characiformes (*S. rhombeus* and *T. albus*). Both sampled clades included one omnivorous and one detritivorous species. Of the omnivorous species, *M. festivus* is mostly detritivorous and feeds off the benthos and from plant detritus (Röpke *et al*., 2013), while *T. albus* feeds at the water surface (diet mostly composed of fruits, seeds and insects) (Gama *et al.,* 2001). Of the carnivorous species, *Cichla spp*. is piscivorous (Aguiar-Santos *et al*., 2018), while *S. rhombeus* will eat anything from fish scales and entire fish to mussels (Sá-Oliveira *et al*., 2017). All four species live on river and lake margins and none of the species shows seasonal migration patterns. All species are often found in large aggregations, although large specimens of *S. rhombeus* and *Cichla spp*. are mostly solitary.

### Supplementary tables

**Suppl. Table 1**: Site identification, water color, geographical coordinates, ecosystem type and sampling time for each sampling site.

| Site characteristics | | | | | | |
|---|---|---|---|---|---|---|
| **Site #** | **Site name** | **Water color** | **GPS S** | **GPS W** | **Ecosystem** | **Sampling time** |
| 1 | Rio Negro - Barcelos | Black | 0°50'50.8"S | 62°57'40.3"W | River | 11/2018 |
| 2 | Rio Negro - Santo Alberto | Black | 1°23'29.8"S | 61°59'35.3"W | River | 10/2019 |
| 3 | Rio Negro - Anavilhanas | Black | 2°41'46.1"S | 60°46'33.3"W | River | 10/2018 |
| 4 | Lago do cemeterio | Black | 3°02'16.6"S | 60°32'42.7"W | Lake | 10/2019 |
| 5 | Lago Téfé | Black | 3°27'55.2"S | 64°53'13.2"W | Lake | 11/2019 |
| 6 | Rio Branco | White | 1°19'05.7"S | 61°52'34.7"W | River | 10/2019 |
| 7 | Lago Janauari | White | 3°12'03.4"S | 60°03'10.1"W | Lake | 10/2018 |
| 8 | Lago Catalão | White | 3°09'56.4"S | 59°54'38.4"W | Lake | 10/2018 |
| 9 | Lago Janauaca | White | 3°23'37.5"S | 60°19'52.6"W | Lake | 11/2018 |
| 10 | Rio Manacapuru | White | 3°16'16.9"S | 60°42'03.2"W | River | 11/2018 |
| 11 | Lago Téfé-Solimões | White | 3°21'07.4"S | 64°40'21.4"W | Lake | 11/2019 |
| 12 | Lago des pirates | White | 3°15'19.2"S | 64°41'44.3"W | Lake | 11/2019 |
| 13 | Balbina Reservoir | Clear | 1°50'55.9"S | 59°34'59.5"W | Reservoir | 10/2018 |
| 14 | Rio Tapajós | Clear | 2°18'57.8"S | 55°00'45.0"W | River | 10/2019 |
| 15 | Rio Curua-Una | Clear | 2°48'19.1"S | 54°17'52.2"W | River | 11/2018 |

**Suppl. Table 2**: Sequencing output of Amazonian fish host gill transcriptomes (NovaSeq).

| Site | Number of reads per species per site | | | |
|---|---|---|---|---|
| | **Flag cichlid** | **Black piranha** | **Sardine** | **Peacock bass** |
| Site #1 | 1.48E+08 | 3.18E+08 | 2.26E+08 | 2.45E+08 |
| Site #2 | 4.28E+08 | 2.81E+08 | 6.07E+08 | 1.86E+08 |
| Site #3 | 3.22E+08 | 3.75E+08 | 4.20E+08 | 3.66E+08 |
| Site #4 | 2.19E+08 | 3.23E+08 | 2.85E+08 | 2.17E+07 |
| Site #5 | 2.99E+08 | 4.35E+08 | 3.19E+08 | 1.78E+08 |
| Site #6 | 3.11E+08 | 2.81E+08 | 3.46E+08 | 7.22E+07 |
| Site #7 | 3.21E+08 | 2.00E+08 | 3.27E+08 | 1.73E+08 |
| Site #8 | 2.97E+08 | 1.97E+08 | 3.06E+08 | 5.71E+07 |
| Site #9 | 1.95E+08 | 1.85E+08 | 3.83E+08 | 3.58E+07 |
| Site #10 | 2.40E+08 | - | - | 1.30E+08 |
| Site #11 | 2.39E+08 | 1.97E+08 | 2.05E+08 | 1.40E+08 |
| Site #12 | 2.84E+08 | 3.66E+08 | 3.44E+08 | 1.52E+08 |
| Site #13 | 1.36E+08 | 1.06E+08 | 2.68E+08 | 9.33E+07 |
| Site #14 | 6.12E+07 | 1.31E+08 | 2.12E+08 | - |
| Site #15 | 1.18E+08 | 2.33E+08 | 1.80E+08 | - |
| Total | 3.62E+09 | 3.63E+09 | 4.43E+09 | 1.85E+09 |
| Global total | 1.35E+10 | | | |

**Suppl. Table 3**: Sequencing output of zebrafish host gill transcriptomes (NovaSeq).

| Water color | Treatment | Reads |
|---|---|---|
| Black water | BWIS | 7.73E+07 |
| | BWIS | 8.25E+07 |
| | BWN | 1.75E+08 |
| | BWS | 1.34E+08 |
| White water | WWIS | 6.52E+07 |
| | WWIS | 1.81E+08 |
| | WWN | 2.28E+08 |
| | WWS | 1.07E+08 |
| Total | | 1.05E+09 |

**Suppl. Table 4**: Number of specimens of each host species collected at each sampling site[1].

| Site # | *M. festivus* | *T. albus* | *S. rhombeus* | *Cichla spp.* |
|---|---|---|---|---|
| 1 | 20 | 20 | 20 | 18 |
| 2 | 20 | 11 | 20 | 17 |
| 3 | 20 | 20 | 20 | 5 |
| 4 | 20 | 0 | 0 | 11 |
| 5 | 20 | 18 | 17 | 4 |
| 6 | 20 | 20 | 20 | 0 |
| 7 | 20 | 16 | 15 | 18 |
| 8 | 20 | 20 | 20 | 20 |
| 9 | 20 | 20 | 20 | 20 |
| 10 | 20 | 20 | 20 | 17 |
| 11 | 20 | 13 | 12 | 0 |
| 12 | 20 | 15 | 15 | 0 |
| 13 | 20 | 9 | 7 | 19 |
| 14 | 20 | 14 | 20 | 13 |
| 15 | 20 | 20 | 20 | 20 |
| Total | 300 | 236 | 246 | 182 |

***1:** "*M. festivus*" stands for *Mesonauta festivus*, "*T. albus*" stands for *Triportheus albus* and "*S. rhombeus*" stands for *Serrasalmus rhombeus*.

**Supp. Table 5**: Measure of DOC quantity and FDOM optical characteristics[1].

| Site # | DOC and FDOM characteristics | | | | | | |
|---|---|---|---|---|---|---|---|
| | DOC conc. | SAC340 | SUVA254 | Abs 254/365 | % humic FDOM | % fulvic FDOM | % protein FDOM |
| 1 | 10.9 | 39.5 | 4.5 | 3.8 | 56.7 | 29.5 | 13.8 |
| 2 | 11.7 | 33.5 | 3.7 | 3.6 | 60.3 | 32.6 | 7.1 |
| 3 | 11.4 | 30.5 | 3.6 | 3.8 | 47.2 | 30.3 | 22.5 |
| 4 | 9.8 | 18.9 | 2.4 | 3.8 | 52.3 | 41.0 | 6.7 |
| 5 | 7.1 | 29.1 | 3.4 | 4.0 | 54.2 | 37.3 | 8.5 |
| 6 | 6.0 | 19.1 | 2.2 | 4.3 | 50.8 | 39.7 | 9.5 |
| 7 | 7.1 | 19.1 | 1.4 | 2.2 | 34.7 | 36.2 | 29.1 |
| 8 | 9.1 | 11.7 | 2.1 | 6.4 | 37.5 | 45.6 | 16.8 |
| 9 | 5.7 | 20.0 | 2.6 | 4.2 | 50.6 | 40.4 | 9.0 |
| 10 | 8.0 | 22.1 | 3.0 | 4.6 | 46.2 | 41.8 | 12.0 |
| 11 | 5.7 | 20.1 | 2.6 | 3.8 | 49.0 | 39.3 | 11.7 |
| 12 | 6.5 | 14.2 | 2.2 | 4.7 | 43.9 | 45.3 | 10.8 |
| 13 | 4.9 | 6.1 | 1.2 | 7.1 | 30.6 | 42.3 | 27.1 |
| 14 | 2.7 | 8.7 | 1.9 | 5.0 | 44.8 | 38.3 | 16.9 |
| 15 | 4.6 | 11.7 | 1.9 | 5.3 | 35.1 | 45.0 | 20.0 |

***1:** “DOC” stands for dissolved organic carbon, “FDOM” stands for fluorescent dissolved organic matter, “DOC conc.” means DOC concentration in mg $L^{-1}$. SAC340 and SUVA254 are the specific absorbance coefficients index of relative DOM aromaticity (higher the values more aromatic is the DOM). Abs254/365 is the index of molecular weight: the lower the value the higher molecular weight is the DOM is.

**Suppl. Table 6**: Concentrations of free ions and nutrients.

| Site # | Water color | Ions: mg $L^{-1}$ | | | | | Nutrients: umol $L^{-1}$ | | |
|---|---|---|---|---|---|---|---|---|---|
| | | $Na^{+}$ | $Mg^{+2}$ | $K^{+}$ | $Ca^{+2}$ | $Cl^{-}$ | Nitrite | Nitrate | Silicate |
| 1 | Black | 0.46 | 0.12 | 0.42 | 0.04 | 0.11 | 0.11 | 3.20 | 64.41 |
| 2 | Black | 0.25 | 0.09 | 0.33 | 0.49 | 1.16 | 0.10 | 2.87 | 92.32 |
| 3 | Black | 1.80 | 0.26 | 0.65 | 0.08 | 0.32 | 0.09 | 4.36 | 72.55 |
| 4 | Black | 0.23 | 0.05 | 0.14 | 0.37 | 0.64 | 0.01 | 0.47 | 76.82 |
| 5 | Black | 0.87 | 0.19 | 0.56 | 0.82 | 0.53 | 0.08 | 4.09 | 217.19 |
| 6 | White | 1.15 | 0.43 | 0.70 | 0.93 | 1.10 | 0.04 | 8.23 | 180.48 |
| 7 | White | 1.99 | 0.20 | 0.79 | 0.06 | 1.47 | 0.19 | 1.31 | 98.31 |
| 8 | White | 4.56 | 3.76 | 1.71 | 0.83 | 1.75 | 0.09 | 0.56 | 242.31 |
| 9 | White | 3.32 | 1.00 | 1.07 | 0.44 | 2.17 | 0.13 | 20.45 | 156.51 |
| 10 | White | 4.91 | 0.14 | 1.45 | 0.05 | 1.43 | 0.12 | 1.53 | 126.01 |
| 11 | White | 1.95 | 0.21 | 0.28 | 1.11 | 1.29 | 0.03 | 6.47 | 326.53 |
| 12 | White | 5.35 | 1.76 | 1.28 | 1.17 | 3.26 | 0.61 | 11.96 | 222.31 |
| 13 | Clear | 0.80 | 0.14 | 0.67 | 0.03 | 0.78 | 0.05 | 1.55 | 85.93 |
| 14 | Clear | 0.43 | 0.47 | 0.57 | 0.68 | 0.39 | 0.09 | 1.91 | 179.36 |
| 15 | Clear | 1.52 | 0.26 | 0.67 | 0.04 | 1.22 | 0.06 | 2.55 | 171.96 |

**Suppl. Table 7**: Primary productivity characterization and measure of physicochemical parameters.[1]

| Site # | Water color | Primary productivity: ug $L^{-1}$ | | | Physicochemical parameters | | | |
|---|---|---|---|---|---|---|---|---|
| | | Chl a | Phaeopig. | Chla/DOC | Temp. °C | Cond. uS | pH | % $O_2$ |
| 1 | Black | 0.35 | 2.43 | 0.03 | 31.60 | 13.10 | 3.71 | 92.12 |
| 2 | Black | 0.73 | 0.33 | 0.06 | 30.60 | 10.60 | 4.16 | 58.00 |
| 3 | Black | 0.05 | 0.38 | 0.00 | 30.70 | 13.20 | 4.24 | 53.20 |
| 4 | Black | 1.35 | 1.44 | 0.14 | 32.40 | 7.20 | 3.83 | 76.90 |
| 5 | Black | 1.82 | 1.73 | 0.26 | 30.00 | 10.60 | 4.98 | 61.50 |
| 6 | White | 6.21 | 2.89 | 1.03 | 31.00 | 22.00 | 6.25 | 88.70 |
| 7 | White | 4.62 | 17.31 | 0.65 | 32.90 | 22.40 | 4.38 | 60.00 |
| 8 | White | 7.14 | 6.60 | 0.79 | 32.90 | 174.80 | 5.70 | 44.00 |
| 9 | White | 1.35 | 1.88 | 0.24 | 29.30 | 88.00 | 6.75 | 82.60 |
| 10 | White | 2.78 | 10.54 | 0.35 | 32.60 | 24.30 | 5.31 | 72.80 |
| 11 | White | 4.41 | 3.20 | 0.77 | 30.30 | 19.70 | 6.05 | 68.60 |
| 12 | White | 9.05 | 4.69 | 1.40 | 31.90 | 127.60 | 7.15 | 31.90 |
| 13 | Clear | 0.83 | 0.78 | 0.17 | 33.20 | 16.80 | 5.05 | 103.20 |
| 14 | Clear | 2.15 | 1.03 | 0.81 | 30.00 | 14.10 | 6.36 | 80.20 |
| 15 | Clear | 1.25 | 2.38 | 0.28 | 31.20 | 19.00 | 6.00 | 79.10 |

***1:** “Chl a” means the concentration of chlorophyll a; “Phaeopig.” means the concentration of phaeopigments; “Chla/DOC” is the ratio of chlorophyll a concentration divided by DOC concentration; “Temp. °C” means the temperature in ° Celsius; “Cond. uS” means the conductivity in microsiemens; “% $O_2$” means the percentage of saturation of dissolved oxygen.

**Suppl. Table 8**: Concentration of dissolved metals in ug/L.

| Site # | Water color | Metals (ug/l) | | | | | | | | | | | |
|---|---|---|---|---|---|---|---|---|---|---|---|---|---|
| | | Al | V | Cr | Mn | Fe | Co | Ni | Cu | Zn | As | Cd | Pb |
| 1 | Black | 137.75 | 0.38 | 0.30 | 7.38 | 166.63 | 0.13 | 1.93 | 10.36 | 33.48 | 0.16 | 0.09 | 1.43 |
| 2 | Black | 150.00 | 0.10 | 0.05 | 5.90 | 160.00 | 0.10 | 0.15 | 0.30 | 11.00 | 0.05 | 0.02 | 0.27 |
| 3 | Black | 36.33 | 0.34 | 0.37 | 9.24 | 142.38 | 0.28 | 3.23 | 9.25 | 72.92 | 0.48 | 0.21 | 1.11 |
| 4 | Black | 87.00 | 0.30 | 0.05 | 4.60 | 100.00 | 0.10 | 0.33 | 1.90 | 9.00 | 0.08 | 0.13 | 0.30 |
| 5 | Black | 62.00 | 0.10 | 0.33 | 13.00 | 220.00 | 0.10 | 0.52 | 0.60 | 4.40 | 0.19 | 0.02 | 0.12 |
| 6 | White | 38.00 | 0.20 | 0.05 | 0.51 | 230.00 | 0.10 | 0.14 | 0.80 | 2.60 | 0.07 | 0.02 | 0.26 |
| 7 | White | 65.50 | 0.78 | 0.40 | 9.85 | 269.28 | 0.10 | 0.85 | 16.19 | 44.15 | 0.47 | 0.06 | 0.67 |
| 8 | White | 1.81 | 0.17 | 0.10 | 0.61 | 5.84 | 0.10 | 0.48 | 2.20 | 171.78 | 0.99 | 0.02 | 0.03 |
| 9 | White | 28.02 | 1.45 | 0.09 | 11.25 | 166.97 | 0.10 | 0.58 | 2.73 | 1.83 | 0.70 | 0.03 | 0.25 |
| 10 | White | 13.47 | 0.85 | 0.21 | 4.64 | 97.85 | 0.10 | 1.12 | 2.11 | 25.85 | 0.38 | 0.08 | 0.16 |
| 11 | White | 49.00 | 0.30 | 0.11 | 0.68 | 250.00 | 0.10 | 0.41 | 0.50 | 2.70 | 0.27 | 0.02 | 0.21 |
| 12 | White | 27.00 | 0.20 | 0.06 | 4.60 | 82.00 | 0.10 | 0.60 | 1.70 | 8.10 | 1.30 | 0.02 | 0.11 |
| 13 | Clear | 10.29 | 0.05 | 0.05 | 0.23 | 16.85 | 0.10 | 0.13 | 0.56 | 4.49 | 0.14 | 0.02 | 0.05 |
| 14 | Clear | 5.00 | 0.10 | 0.05 | 0.05 | 7.00 | 0.10 | 0.10 | 0.50 | 3.70 | 0.07 | 0.02 | 0.03 |
| 15 | Clear | 18.49 | 0.17 | 0.58 | 12.31 | 52.88 | 0.11 | 1.18 | 2.12 | 23.94 | 0.64 | 0.07 | 0.25 |

**Suppl. Table 9:** Comparison of water parameters measured in the original sample and after transport to the laboratory (prior to the axenic zebrafish experiment).[1]

| Parameter | Measure | Blackwater Santo Alberto | | Whitewater Lago des pirates | |
|---|---|---|---|---|---|
| | | Original sample | After transport | Original sample | After transport |
| [Al] | µg/L | 150.00 | 113.00 | 27.00 | 11.00 |
| [As] | µg/L | <0.30 | <0.30 | 1.30 | 0.90 |
| [Cd] | µg/L | <0.10 | <0.10 | <0.10 | <0.10 |
| [$Ca^{+2}$] | µg/L | 490.00 | 127.00 | 1170.00 | 1530.00 |
| [Cr] | µg/L | <0.50 | <0.50 | <0.50 | <0.50 |
| [Co] | µg/L | <0.50 | <0.50 | <0.50 | <0.50 |
| [Cu] | µg/L | 0.30 | 12.60 | 1.70 | 9.40 |
| [Fe] | µg/L | 160.00 | 175.00 | 82.00 | 38.00 |
| [$Mg^{+2}$] | µg/L | 90.00 | 55.00 | 1176.00 | 2320.00 |
| [Mn] | µg/L | 5.90 | 4.00 | 4.60 | 1.00 |
| [Ni] | µg/L | <1.00 | <1.00 | <1.00 | <1.00 |
| [Pb] | µg/L | 0.27 | <0.10 | 0.11 | <0.10 |
| [$K^{+}$] | µg/L | <500.00 | <500.00 | 1280.00 | 1300.00 |
| [$Na^{+}$] | µg/L | 250.00 | 430.00 | 5350.00 | 4490.00 |
| [V] | µg/L | <0.50 | <0.50 | <0.50 | <0.50 |
| [Zn] | µg/L | 11.00 | 4.00 | 8.10 | <3.00 |
| [DOC] | mg/L | 11.67 | 10.40 | 6.47 | 5.84 |
| Conductivity | µS | 11.00 | 22.00 | 128.00 | 128.00 |
| pH | - | 4.16 | 4.05 | 7.15 | 7.00 |

***1:** “DOC” stands for dissolved organic carbon.

**Suppl. Table 10**: Proteins known to be involved in ionoregulatory processes in fish facing acidic and ion-poor environments.[1]

| Protein | Strategy | Reference |
|---|---|---|
| $Na^+/H^+$ antiporters (NHE) | Strategy #2 | Morris et al. 2021 |
| V-type proton ATPases | Strategy #2 | Morris et al. 2021 |
| Sodium channels | Strategy #2 | Morris et al. 2021 |
| Rh glycoproteins (Rhag, Rhbg, Rhcg) | Strategy #2 | Morris et al. 2021 |
| Carbonic anhydrases | Strategy #2 | Morris et al. 2021 |
| $Na^+/K^+$ ATPases | Strategy #2 | Morris et al. 2021 |
| Sodium/Potassium/Calcium exchangers | Strategy #2 | Morris et al. 2021 |
| Electrogenic $Na^+/HCO3^-$ cotransporters | Strategy #2 | Morris et al. 2021 |
| $Cl^-/HCO_3^-$ exchangers | Strategy #2 | Guh et al. 2015 |
| Anion exchangers (e.g. $Cl^-$ exchanger) | Strategy #2 | Guh et al. 2015 |
| Chloride channels | Strategy #2 | Guh et al. 2015 |
| Calcium channels | Strategy #2 | Guh et al. 2015 |
| Potassium channels | Strategy #2 | Guh et al. 2015 |
| $Na^+$ $Cl^-$ cotransporters | Strategy #2 | Guh et al. 2015 |
| $Na^+$ $Ca_2^+$ exchangers | Strategy #2 | Guh et al. 2015 |
| $Ca_2^+$ ATPases | Strategy #2 | Guh et al. 2015 |
| Solute carriers (SLC) | Strategy #2 | Guh et al. 2015 |
| Occludins | Strategy #1 | Araujo et al. 2017 |
| Claudins | Strategy #1 | Araujo et al. 2017 |
| Actinin | Strategy #1 | Araujo et al. 2017 |
| Integrins | Strategy #1 | Araujo et al. 2017 |
| Desmoplakins | Strategy #1 | Araujo et al. 2017 |
| Gap junction proteins | Strategy #1 | Araujo et al. 2017 |
| Glucocorticoid receptors | Strategy #1 | Araujo et al. 2017 |
| Prolactin receptors | Strategy #1 | Araujo et al. 2017 |
| Mineralocortocoid receptors | Strategy #1 | Chasiotis et al. 2012 |
| Zonula occludens (plaque proteins) | Strategy #1 | Chasiotis et al. 2012 |
| Protein kinase C | Strategy #1 | Ulluwishewa et al. 2011 |
| Mitogen-activated protein kinase | Strategy #1 | Ulluwishewa et al. 2011 |
| Myosin light chain kinase | Strategy #1 | Ulluwishewa et al. 2011 |
| Rho family of small GTPases (RhoA, Rac, Cdc42, etc) | Strategy #1 | Ulluwishewa et al. 2011 |
| Rho kinases | Strategy #1 | Ulluwishewa et al. 2011 |
| Tricellulins | Strategy #1 | Ulluwishewa et al. 2011 |
| JAM proteins (JAM-A, coxsackie/adenovirus receptor) | Strategy #1 | Ulluwishewa et al. 2011 |
| Toll-like receptors | Strategy #1 | Ulluwishewa et al. 2011 |

| | | |
|---|---|---|
| Cadherins | Strategy #1 | Ghosh et al. 2021 |
| Catenins | Strategy #1 | Ghosh et al. 2021 |
| Desmogleins | Strategy #1 | Ghosh et al. 2021 |
| Desmocollins | Strategy #1 | Ghosh et al. 2021 |
| Connexins | Strategy #1 | Ghosh et al. 2021 |

***1:** Strategy #1 refers to the regulation of ion efflux and strategy #2 refers to the regulation of active ion uptake.

### Supplementary figures

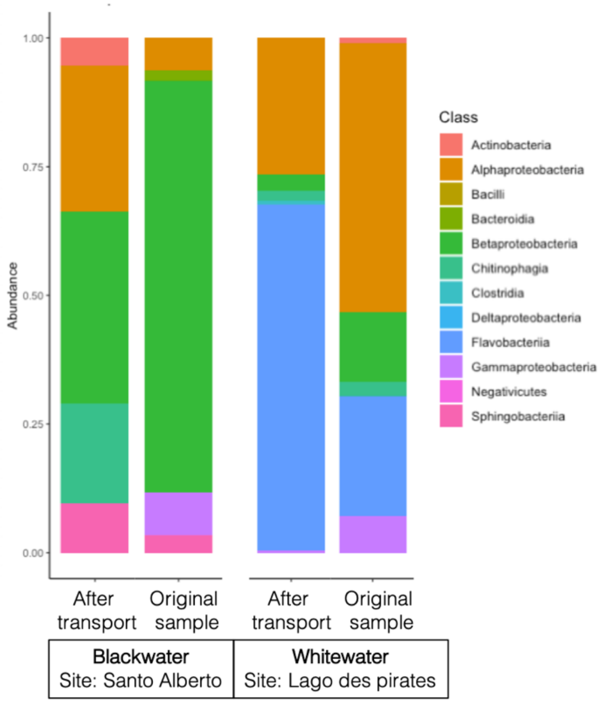


**Suppl. Fig. 1**: Stacked barplot of the relative abundance of the bacterial classes detected in water samples used for the axenic zebrafish experiment (16S rRNA dataset). The "Original sample" was taken directly on site, and the "After transport" sample was taken after transportation of the water from the field to the laboratory, right before the start of the axenic zebrafish experiment. In the blackwater from the Santo Alberto site, the sample taken after transportation shows a decrease in Betaproteobacteria and Gammaproteobacteria, and an increase in Actinobacteria, Alphaproteobacteria and Chitinophagia. In the whitewater from Lago des pirates, the sample taken after transportation shows an increase in Flavobacteriia, and a decrease in Proteobacteria (Alphaproteobacteria, Betaproteobacteria and Gammaproteobacteria).

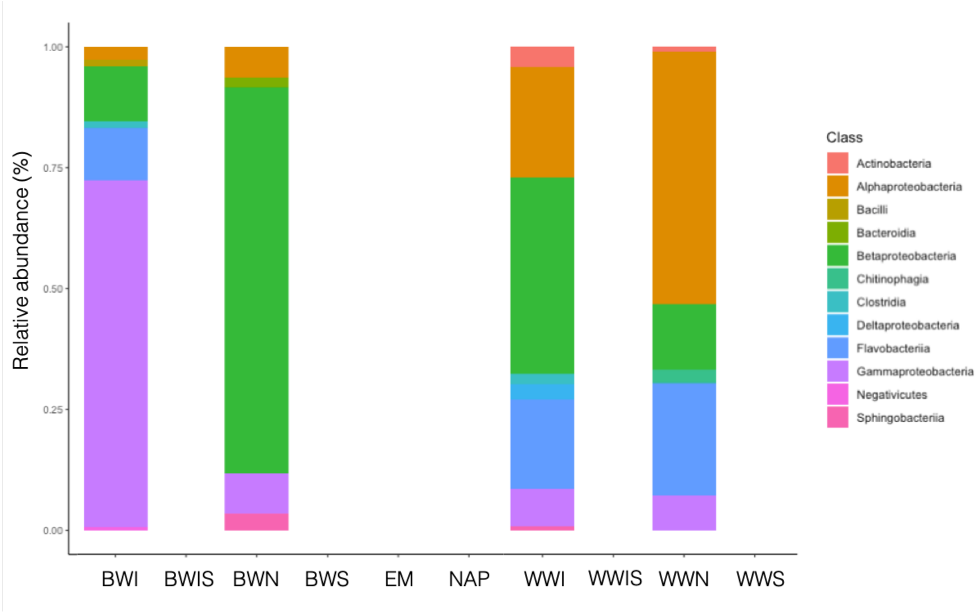


**Suppl. Fig. 2**: Stacked barplot of the relative abundance of the bacterial classes detected in the **water samples** collected during the axenic zebrafish experiment (16S rRNA dataset). "BWI" stands for inverted non-sterile blackwater, "BWIS" for inverted sterile blackwater, "BWN" for non-sterile blackwater, "BWS" for sterile blackwater, "EM" for embryo medium (the medium in which zebrafish larvae were kept for the first 96h), "NAP" for nucleic acid preservation buffer (to control potential cross-contamination from the buffer), "WWI" for inverted non-sterile whitewater, "WWIS" for inverted sterile whitewater, "WWN" for non-sterile whitewater, and "WWS" stands for sterile whitewater. Overall, this plot shows that after filtration of potential contaminant sequences using the decontam R package (Davis *et al.,* 2018), and after removal of sequences appearing <2 times in at least one of the experimental groups, we do not detect any ASV in sterile and in control groups.

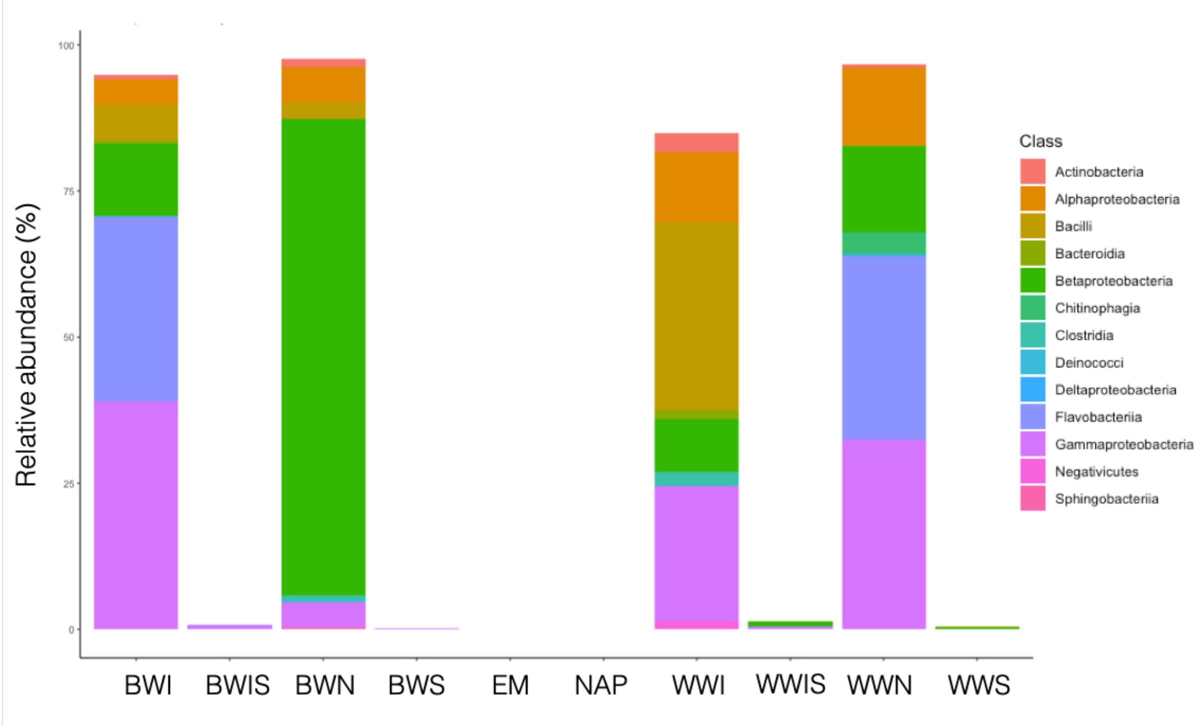


**Suppl. Fig. 3**: Stacked barplot of the relative abundance of the bacterial classes detected in the **zebrafish larvae samples** collected during the axenic zebrafish experiment (all groups in this plot except "EM" and "NAP" are fish samples) (16S rRNA dataset). "BWI" stands for inverted non-sterile blackwater zebrafish, "BWIS" for inverted sterile blackwater zebrafish, "BWN" for non-sterile blackwater zebrafish, "BWS" for sterile blackwater zebrafish, "EM" for embryo medium (the medium in which zebrafish larvae were kept for the first 96h, shown here as control), "NAP" for nucleic acid preservation buffer (to control potential cross-contamination from the buffer, shown here as control), "WWI" for inverted non-sterile whitewater zebrafish, "WWIS" for inverted sterile whitewater zebrafish, "WWN" for non-sterile whitewater zebrafish, and "WWS" stands for sterile whitewater zebrafish. Overall, this plot shows that after filtration of potential contaminant sequences using the decontam R package (Davis *et al*., 2018), and after removal of sequences appearing <2 times in at least one of the experimental groups, we detect a very low number of ASVs in sterile and in control groups.

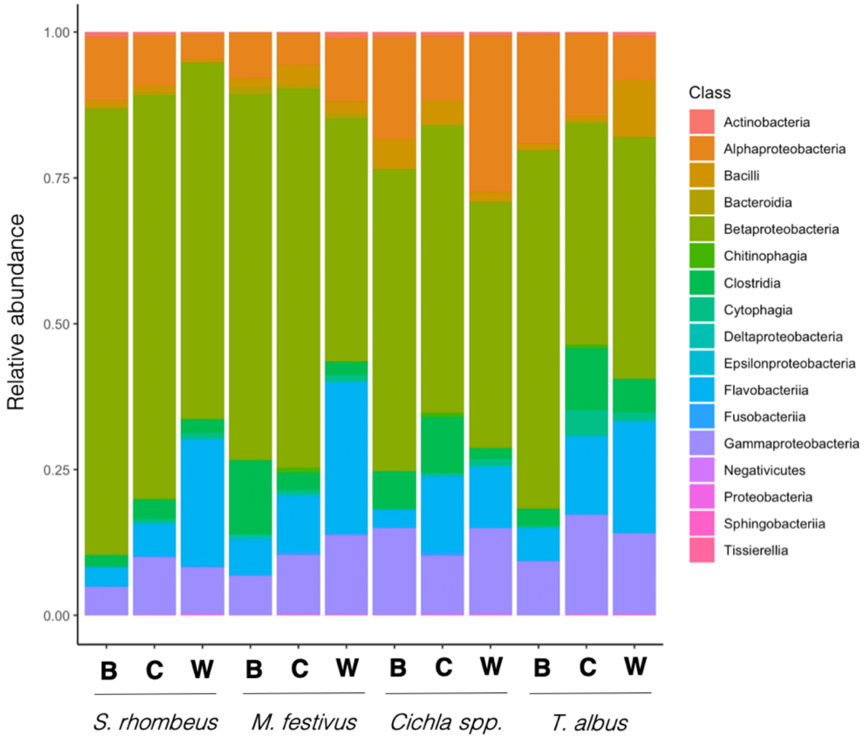


**Suppl. Fig. 4:** Stacked barplot of the relative abundance of the bacterial classes detected in the Amazonian fish gills microbiomes (16S rRNA gene dataset). "*M. festivus*" stands for *Mesonauta festivus*, "*T. albus*" stands for *Triportheus albus* and "*S. rhombeus*" stands for *Serrasalmus rhombeus*. The letters in bold indicate water type: "B" for blackwater, "C" for clearwater and "W" for whitewater.

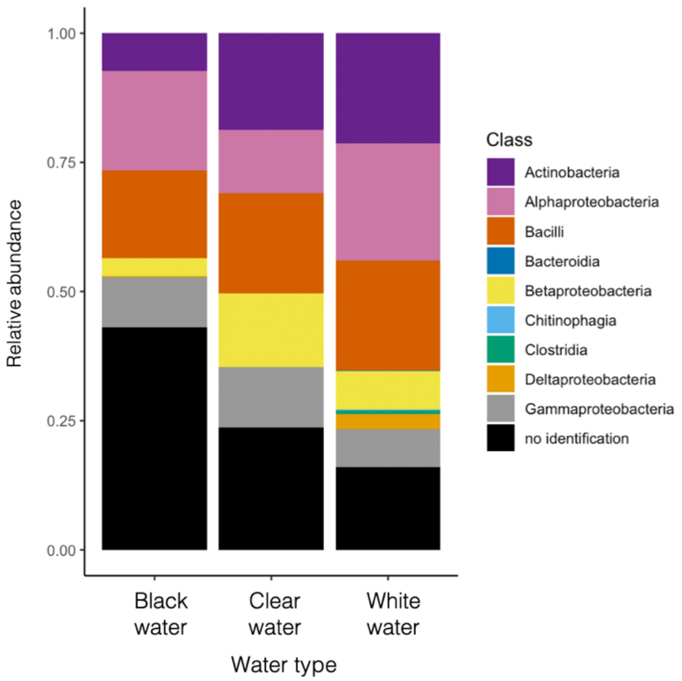


**Suppl. Fig. 5:** Stacked barplot of the relative abundance of the bacterial classes detected in the bacterioplankton samples collected at the 15 Amazonian field sampling sites (16S rRNA gene dataset).

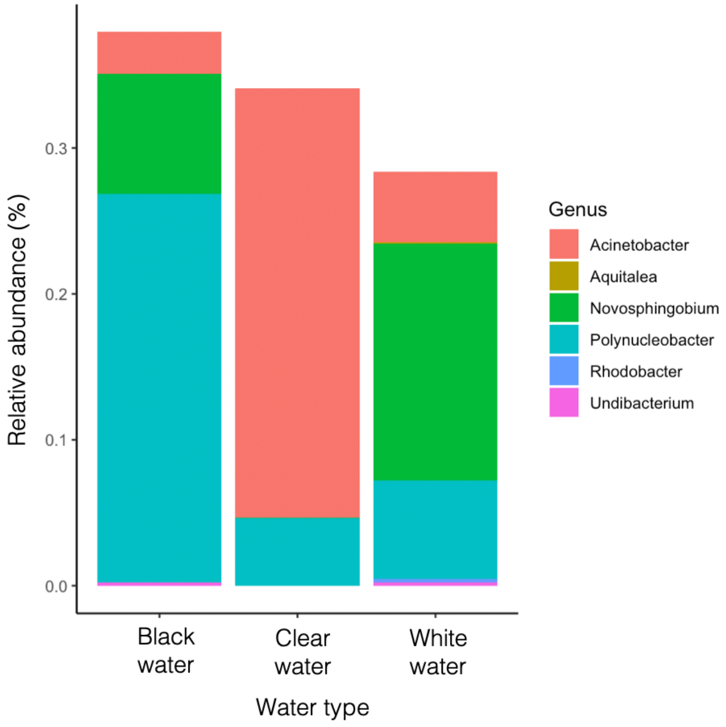


**Suppl. Fig. 6:** Stacked barplot showing the relative abundance of the fish gill bacterial biomarkers in the free-living bacterioplanktonic communities collected at the 15 Amazonian field sampling sites (16S rRNA gene dataset). The bacterial biomarkers not shown were not detected in the bacterioplankton.
